# The psychedelic, DOI, increases dopamine release in nucleus accumbens to predictable rewards and reward cues

**DOI:** 10.1101/2024.03.29.587390

**Authors:** David Martin, Angel Delgado, Donna Calu

## Abstract

Psychedelics produce lasting therapeutic responses in neuropsychiatric diseases suggesting they may disrupt entrenched associations and catalyze learning. Here, we examine psychedelic effects on dopamine signaling in the nucleus accumbens (NAc) core, a region extensively linked to reward learning, motivation, and drug-seeking. We measure phasic dopamine transients following acute psychedelic administration during well learned Pavlovian tasks in which sequential cues predict rewards. We find that the psychedelic 5-HT_2A/2C_ agonist, DOI, increases dopamine signaling to rewards and proximal reward cues but not to the distal cues that predict these events. We determine that the elevated dopamine produced by psychedelics to reward cues occurs independently of psychedelic-induced changes in reward value. The increased dopamine associated with predictable reward cues supports psychedelic-induced increases in prediction error signaling. These findings lay a foundation for developing psychedelic strategies aimed at engaging error-driven learning mechanisms to disrupt entrenched associations or produce new associations.

## Introduction

Psychedelic 5-HT_2A_ agonists including psilocybin, LSD, and 2,5-dimethoxy-4-iodoamphetamine (DOI) produce profound acute subjective effects and accumulating evidence indicates psychedelics have significant clinical utility in the treatment of neuropsychiatric conditions including anxiety, depression, and addiction[1–5]. Acute administration of psychedelic drugs can engender positive alterations in mood and attitudes that persist for months[6,7]. These changes suggest that psychedelic effects on reward learning pathways may be important therapeutic mechanisms. Here, we aimed to study the effects of psychedelics on reward prediction error signaling, a fundamental element of associative learning, to gain insight into psychedelic mechanisms.

Ventral tegmental dopaminergic neuronal activity and phasic dopamine (DA) release in the nucleus accumbens core (NAc) are strongly implicated in reward prediction error signaling[8–11], learning[12], and drug-seeking[13,14]. DA is phasically released in NAc in response to unpredictable reward delivery, but as learning progresses, DA release is inhibited to reward and shifted to the earliest stimuli predictive of reward[8,10]. DA promotes the formation of conditioned behaviors[15,16] and supports associative model-based learning[17], highlighting its role as a ‘teaching’ signal in the context of reward prediction error based models of DA function[18]. Other roles for DA in NAc include encoding value[19], motivation[20], and salience[21,22].

Psychedelics have been proposed to affect prediction error signaling[23,24]. In humans, low-dose LSD impacts reward-related EEG activity potentially related to increased RPE processing[25], and moderate doses of LSD increase reinforcement learning rates, consistent with greater sensitivity to reward prediction errors (RPEs)[26]. Tonic DA levels are increased by psychedelics in the striatum in humans [27] and in rodents[28], however, measuring psychedelic impacts on fast, phasic DA transients is necessary to determine if psychedelics alter DA-mediated RPE signaling.

To determine the effect of psychedelic 5-HT_2A/2C_ agonist, DOI, on DA signaling during reward prediction we use fiber photometry to measure optical dopamine sensor GRAB_DA_ fluorescence in the NAc during Pavlovian tasks. Because psychedelics alter reward value[29,30], parsing the influence of stimuli value versus stimuli predictability is important for interpreting effects on phasic DA. To achieve this, we utilize two reward types (water and food) in our studies to infer the influences of value and predictability on NAc DA signaling.

## Methods

### Experimental subjects

All subjects were wildtype Sprague-Dawley rats (Charles River). We used female (n=4) and male (n=4) rats across two cohorts for Pavlovian experiments measuring dopamine. For the water behavioral economics experiment we used 8 rats (n=5 male; n=3 female), and for the food behavioral economics experiment we used 14 rats (n=7 male; n=7 female). Rats weighed between 215-375g before the start of the experiments. Rats were single-housed and maintained on a 12:12h reverse light-dark cycle (ZT0 at 1900) and experiments were performed during the dark cycle. For conditions using water reward, rats were water restricted ∼23 hours prior to the experiment and given 15 minutes of water access each day following behavioral experiments. For conditions using food reward, rats were mildly food restricted to maintain a stable weight of ∼95% free feeding weight and fed immediately after behavioral experiments. Free access to water was available over the weekends, and rats were run 4-5 sessions/week. All procedures were performed in accordance with the “Guide for the care and use of laboratory animals” (8th edition, 2011, US National Research Council) and were approved by the University of Maryland School of Medicine Institutional Animal Care and Use Committee (IACUC).

### Surgical Procedures

We anesthetized rats by induction with 5% isoflurane (VetOne). We gave a subcutaneous injection of the analgesic, carprofen (5mg/kg) and a subdermal injection of lidocaine (10mg/ml; ∼0.1 mls) at the incision site prior to the first incision. We infused 600 nL of AAV9-hSyn-GRAB_DA2m[31] (GRAB_DA_) targeting the Nucleus Accumbens Core unilaterally at a 6-degree angle (coordinates from bregma: +1.8 mm AP, +2.15 mm ML, -6.6 mm DV). Following virus infusion, we implanted one 400 μm core, 0.50 NA fiber optic (Thor labs) in the NAc, 0.1mm dorsal to the virus injection coordinates. Fiber optics were first cemented in place with Metabond (Parkell Inc., New York, New York), then with Denmat. The entire headcap was covered in a thin layer of dental cement. Rats were given 3-5 weeks of recovery before testing. All coordinates given are distance from bregma according to the Paxinos and Watson rat brain atlas (6th edition, 2007).

### Histology

At the end of experiments, rats were deeply anesthetized with isoflurane and transcardially perfused with 100 ml of 0.1M sodium phosphate buffer (PBS), followed by 400 ml of 4% paraformaldehyde (PFA) in PBS. Brains were postfixed in 4% PFA for 24 hours before we transferred them to 30% sucrose in PBS for 48 -72h at 4 °C. We stored brains at −20 °C until sectioning. Coronal sections (40 μm) containing NAc core were collected using a cryostat (Leica Microsystems). To verify fiber optic cannula placements, we mounted and coverslipped all brain sections with Vectashield DAPI (Vector Laboratories, Burlingame, CA) and the mounting medium, Mowiol (Sigma-Aldrich, St. Louis, MO). We used a spinning disk confocal microscope (Leica SP8) to verify viral expression using native GRAB_DA_ fluorescence and fiber optic/cannula placement. Rats were excluded if there was no virus expression below the fiber optic in the NAc or if placements were not in the NAc core.

### Behavioral apparatus

Experiments were conducted in individual sound-isolated standard experimental chambers (25 × 27× 30cm; Med Associates). Each chamber had one white house light (6 W) located at the top of a wall that was illuminated for the duration of each session. Food reward conditions used a recessed foodcup and a programmed pellet dispenser that delivered 45 mg food pellets (catalog#1811155; Test Diet 5TUL; protein 20.6%, fat 12.7%, carbohydrate 66.7%). One retractable lever was positioned on either side of the food and water dish, counterbalanced, 6cm above the floor. A speaker producing a tone and another producing white noise were placed in the front top corners of the box. For water reward conditions, an audible syringe pump delivered water. All pokes (entries and exits) into the foodcup/water port were recorded by a photobeam sensor in front of the port.

#### Fiber photometry

Rats were connected via a fiber optic cable (1 m long; 400 μm core; 0.48 NA; Doric) running from the behavioral box to the photometry rig. We used LEDs (ThorLabs) to deliver 465 nm light to measure GRAB_DA_ fluorescence signals and 405 nm light as an isosbestic control. The two wavelengths of light were sinusoidally modulated at 210 and 337Hz respectively. The LEDs connected to a fluorescence mini cube (Doric Lenses). The combined LED output passed through the fiber optic cable which was connected to implanted fiber optics. LED light collected from the GRAB_DA_ and isosbestic channels was focused onto a femtowatt photoreceiver (Newport). We low-pass filtered and digitized the emission light at 3Hz and 5 KHz respectively by a digital processor controlled by Synapse software suite (RZ5P, Tucker Davis Technologies (TDT)). We time-stamped all relevant behavioral data through TTL pulses from MedPC system to Synapse software. Photometry data was sampled at 1017 Hz.

### Behavioral Training

The behavioral economics task used a within-subject design wherein fixed ratios (FRs) were increased across the 30 min session comprised of 10 three-minute bins (FR6, FR10, FR16, FR25 X 2, FR40 X 2, FR63 X 3). Operandi were levers and completion of FRs resulted in activation of syringe pumps delivering 95 uL of water over 1.5 s or the delivery of (2) 45 mg food pellets.

For the experiments using GRAB_DA_ to measure dopamine, rats were trained and recorded during 14-18 Pavlovian training sessions with randomly intermixed 15 CS+ and 15 CS-trials with a 40-60 s inter-trial interval (ITI). The CS+ (tone/white noise, counterbalanced) sounded for 5 s, terminating simultaneously with audible syringe pump activation for 1.5 s to deliver water (95 ul). CS-trials used the opposite audio stimuli (tone or white noise), which signaled for 5 s with no reward delivery.

Following initial Pavlovian training sessions, rats were sequentially tested under a variety of conditions using different cue types (auditory, levers), rewards (food, water), and timing to elucidate the effects of psychedelics on DA using diverse parameters, and selected results are presented here. The exact behavioral history of rats between the 2 cohorts was not identical, though all rats had similar amounts and types of Pavlovian training history, including similar amounts of DOI experience during food and water training prior to collecting the data shown here.

For test sessions with water reward and auditory cues (Figure 3), the CS (distal cue) was 2 s, followed by a gap of 2 s, followed by activation of the audible syringe pump for 1.5 s (95 ul water per delivery). Sessions had 12 ‘Expected’ trials in which the CS preceded the reward intermixed pseudorandomly with 12 ‘Unexpected’ trials in which there was no CS. Test sessions with lever cues (Figure 4) had 15 trials where a lever was extended for 3 s, followed by a 2 s gap, followed by another 3 s lever extension and another 2s gap before proximal cue and reward delivery. In water experiments sequential levers were followed by syringe pump activation for 1.5 s (95 ul) and in food experiments (2) food pellets were delivered.

#### Test Sessions using DOI

For each of the behavioral conditions under study, rats were trained until their behavior and dopamine signals stabilized before being tested in a within-subject counterbalanced design in which each subject received DOI or saline on alternate test days, separated by a retraining session in which no injections were given. For behavioral economic tasks, DOI (0.8 mg/kg i.p) or vehicle (saline) were given 25 min prior to test sessions that were identical to training sessions. In test sessions measuring GRAB_DA_ using audible cues and water reward (Figure 3), 3 cumulative DOI doses were given as a series of 3 injections (400 ug/kg, i.p. for each injection), each before three closely spaced identical 15 min test sessions, with a gap of ∼2 min between sessions to give injections. The 1^st^ injection was given in homecage 15 min prior to 1^st^ session. In test sessions using lever cues (Figure 4), 2 cumulative DOI doses were given as a series of 2 injections (500 ug/kg, i.p. for each injection) immediately prior to each identical 15 min test session with ∼ 2 min between sessions. We note that the peak effect of these cumulative DOI doses is likely not reached until later in the sessions due to lag between intraperitoneal injections and peak drug effects. Nevertheless, this design permits clear observation of dose-response relationships between cumulative DOI doses and DA signals and behavior. All injections were given i.p. at a volume corresponding to 1ml/kg body weight. 2,5-Dimethoxy-4-iodoamphetamine (DOI, Cayman) was dissolved in sterile saline vehicle.

### Data and Statistical Analyses

#### Photometry signal analysis

Raw data were unpacked in Matlab using scripts provided by TDT. Photometry signals were snipped into discrete segments around each trial in the session and F465 signals were fit to F405 (isobestic) signals across all trials using a least-squares linear fit to detrend data. The fitted F405 values for each timepoint were subtracted from the corresponding F465 values, providing a signal for each trial adjusted by the isobestic control[32]. These trial signals were z-scored based on a 5 second baseline period prior to the CS (distal cue) by subtracting the mean during the baseline and dividing by the standard deviation during the baseline. Z-scored traces from all trials of a particular type were averaged within a subject to create an average for that subject and trial type to be used in further analysis. Rats that did not have mean photometry signals significantly greater than baseline (95% confidence interval>0) at the distal cue during their saline test sessions were excluded from analysis. Trials in which signal was interrupted (patch cable disconnection) were manually removed from the data.

Across all experiments, we plot average signal traces with standard error of the mean along with time bins where significance was reached indicated just above the x-axis. Significance across all bins in the trial was determined by calculating 95% confidence intervals (95% CIs) across trial lengths. For within subject comparisons across drug and control conditions, CIs were calculated by subtracting at each time bin the control signal from the drug signal for each individual subject, and 95% CIs were calculated on the group means of these within-subject differences. Differences between control and drug conditions were considered significant when 95% CIs did not contain 0 for at least 350 consecutive milliseconds. This ‘minimum period of significance’ limits false positives arising during brief periods. However, during specific time windows following the onset of cues (from cue onset through 1 second following cue offset) - when there was an *a priori* hypothesis that the drug may have a significant effect – no filter for a minimum period of significance was used.

To summarize data at important timepoints in trials across doses and reinforcers, we measured average GRAB_DA_ signal peak heights (in terms of z-score) during the 1 second period following cues, and/or area under the curve (integrating z-score over time relative to baseline) for the 3 seconds following reward delivery (pump offset/ 1^st^ pellet delivery). We analyzed this data using standard repeated measures, 2-Way ANOVAs using within subject factors across dose, reinforcer, and cue-type where appropriate. Sphericity was not assumed, and Geisser-Greenhouse corrections were employed when required. Post-hoc comparisons between groups were corrected with Dunnett’s multiple testing procedure using Prism software (GraphPad).

#### Behavioral analysis

Data were analyzed in Matlab. In dopamine experiments, reward well poke onset times and offset times were recorded along with lever press onset and offset times. The average probability of a leverpress or reward poke occurring during all timepoints across a trial was calculated for each rat. For within-subject comparisons of drug and control conditions, confidence intervals were calculated by subtracting the control from the drug probabilities for each subject. Periods where the 95% CI does not contain zero are labeled as significant on poking traces with no minimum period of significance. We summarized latency to enter reward well following cues across experiments and these data were analyzed with standard RM ANOVAs and multiple testing correction procedures for post-hoc tests (Dunnett’s).

Behavioral economic analysis was performed by fitting the rewards earned at each price (consecutive bins with the same price were averaged) to the exponential behavioral economics equation[33] for each individual using the “fitnlm” function in matlab. From this data, we extracted best fit values for alpha and Q_0_ for all rats using a fixed value of k=1.2. These data were compared between food and water rewards using standard 2-Way ANOVAs. We also analyzed the number of rewards earned at each price during drug and control conditions using 2-Way ANOVA. Post-hoc comparisons between groups were corrected with Sidak’s multiple testing procedure using Prism software (GraphPad).

Simple linear regression between behavior and GRAB_DA_ signals was performed on training data between each subjects’ average GRAB_DA_ peak signal to the distal cue and the time they spent in the well during the distal cue. To capture relative changes in behavior and DA across time, each subject’s data was normalized across trial blocks such that the minimum and maximum values across trial blocks were set to 0 and 1, respectively, for both poke and signal data. Simple linear regression was also performed between changes in proximal cue GRAB_DA_ peak signals and changes in latency to enter the reward well during DOI experiments on an individual session basis. Changes in GRAB_DA_ signal peak heights to proximal cues between drug and control sessions were normalized to the size of distal cue peaks during control sessions. For these analyses: ΔLatency = Latency_DOI_ – Latency_CON_. ΔDA = (DA_DOI,Proximal_ – DA_CON,Proximal_)/DA_CON,Distal._ All correlations were calculated in Prism.

## Results

### DOI bidirectionally modulates water and food value

NAc DA release is influenced by both reward value and reward predictability[34]. We aimed to design an experiment to disambiguate motivational value and reward prediction in order to interpret psychedelic induced changes in DA signaling relating to these two factors. To do so, we first identified two reward types that show distinct motivational effects in the psychedelic state. We tested the effects of DOI on instrumental lever responding for either water or food reward in a behavioral economics task in which fixed ratio (FR) requirements increase across successive bins of the session. Figure 1A and 1B show behavioral economic curves for food (Fig.1A) and water (Fig.1B) comparing saline and DOI (0.8 mg/kg, i.p.) conditions. We fit the exponential behavioral economic equation[33] to each subject’s curves, revealing an interaction between Reward type and Treatment on consumption at low cost (Q_0_ values: F(1, 20) = 7.429, p=0.013, 2-Way ANOVA, Fig. 1C). Paired t-tests of Q_0_ values indicate that DOI decreases Q_0_ for food (p=0.041) but increases Q_0_ for water (p=0.0042). These data show that motivation for food and water reward at low prices is bidirectionally modulated by administration of DOI. Consistent with opposite effects of DOI on Q_0_ for food and water, at low work requirements (FR6), DOI (0.8 mg/kg, ip.) decreases food consumption, but increases water consumption (Fig 1A,B; Food: t(13)=4.676; p=0.0026; Water: (t(7)=4.968, p=0.0097; Sidak’s multiple comparison correction). At higher prices (FR40, FR63), DOI decreases water consumption (t(7)> 5.487, p’s<0.006, Sidak’s, Fig 1A). DOI also decreases food consumption significantly at FR10, FR16, and FR25 (t(13)>3.689, ps<0.0082, Sidak’s, Fig 1B). Consistent with similar effects of DOI at high costs for both food and water, DOI increases economic demand elasticity (α), or the rate at which consumption decreases with increasing cost, for both water and food reward (main effect of Drug, F(1, 20)=16.37, p=0.0006, Figure 1D).

**Figure 1:**
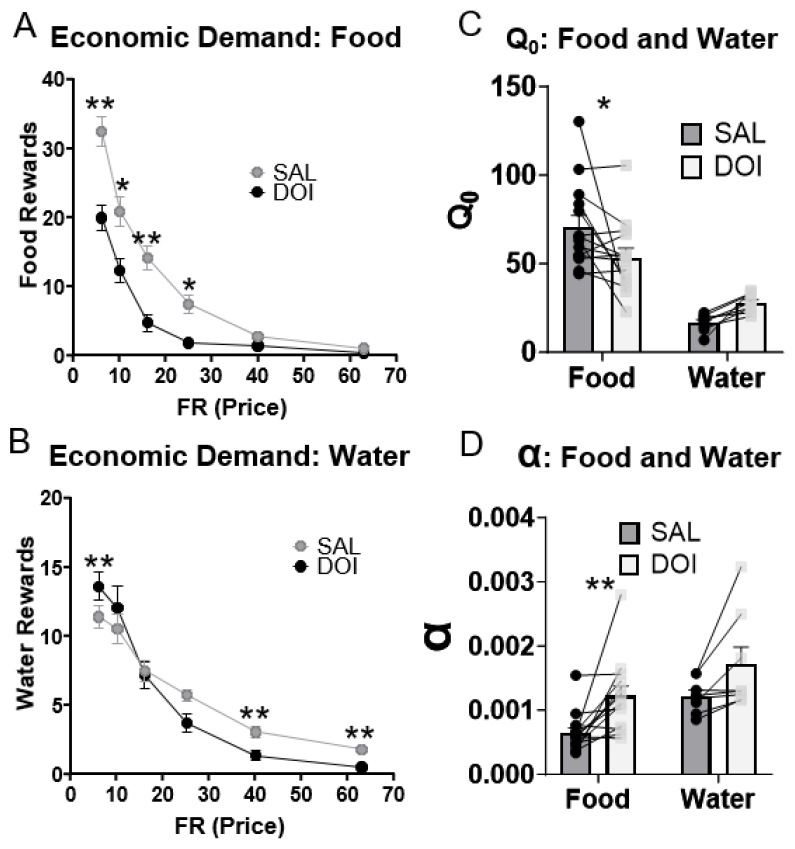
DOI produces bidirectional effects on food and water consumption at low prices: Economic demand data for animals treated with either saline or DOI (0.8 mg/kg, i.p.) prior to behavioral economic testing for water (N=8) or food (N=14). A) Food consumption. B) Water consumption C) Fitted Q_0_ values for food and water. 2 Way AVOVA indicates an interaction between drug and reward type (F(1,20)=7 43; p=0 013). D) Fitted alpha values for food and water. *All Graphs*: “*” “**” indicates p<0.05, p<0.01 at indicated post-hoc comparisons using two way ANOVA and Sidak multiple testing correction.

In the following experiments, we examine the effects of DOI on NAc DA dynamics in Pavlovian conditioning tasks, in which rewards are earned with very low effort - by merely approaching the reward when cues signal their availability. By identifying rewards for which motivation is bidirectionally altered in the psychedelic state, we are positioned to interpret NAc DA signal changes related to factors of predictability and value of rewards and reward cues in subsequent experiments.

### Training shifts NAc dopamine release to distal cues

We infused optical dopamine sensors (GRAB_DA_) and implanted an optical fiber targeting the nucleus accumbens core (NAc) in 8 rats (Figure 2A,B). We water deprived and trained the rats on a Pavlovian task (Figure 2C) in which water reward delivered by an audible syringe pump (proximal cue) was preceded by an audible CS+ (distal cue, tone or white noise) for 5 seconds.

**Figure 2:**
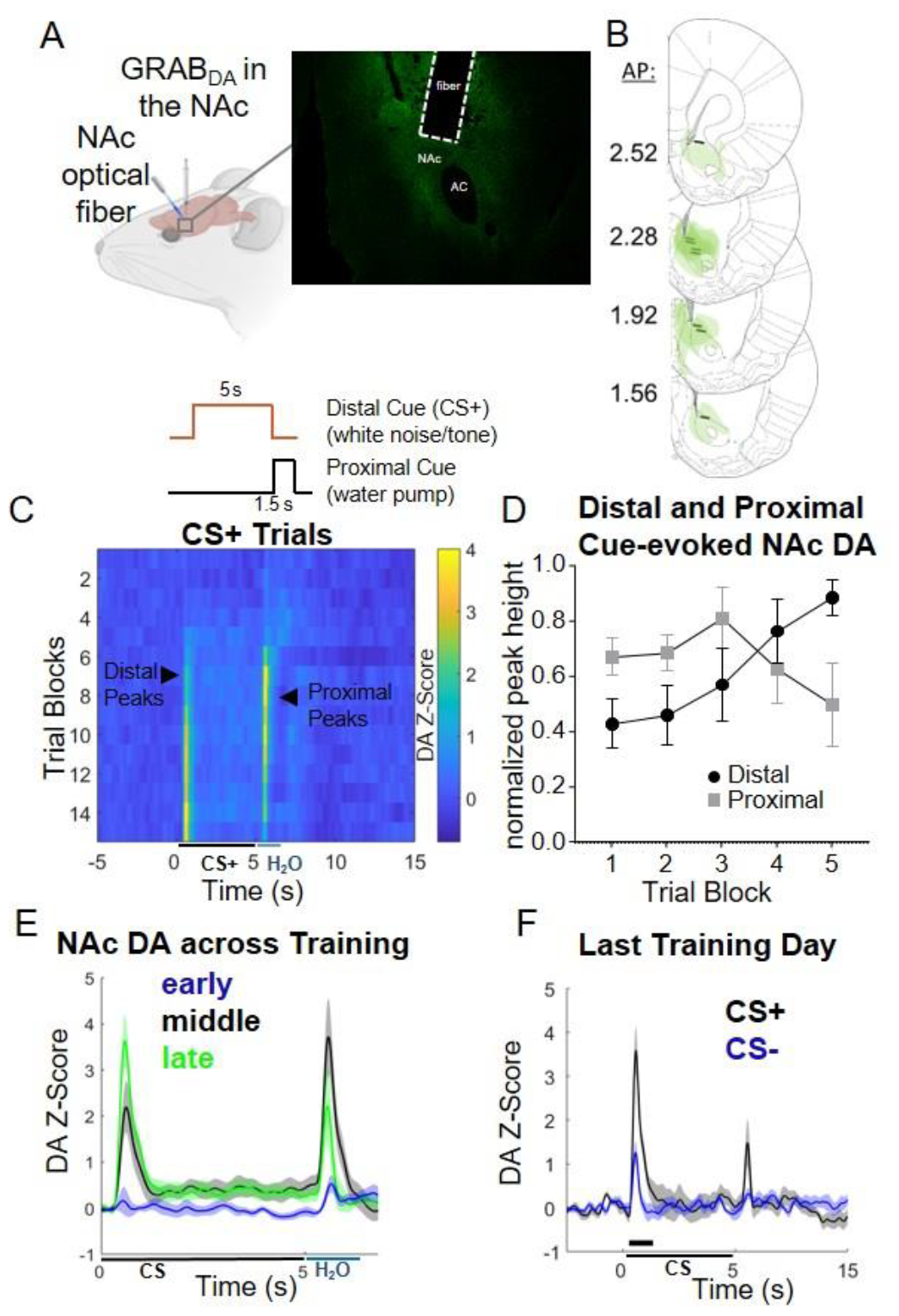
GRAB_DA_ measurement of NAc dopamine during Pavlovian Training: A) Surgical Procedures (left), representative fiber placement (right) B) Fiber tip placements (black bars) and GRAB_DA_ expression (green shading) C) Behavioral Paradigm for training (top), and heat plots of GRAB_DA_ traces across training. CS+ trials are binned into 15 blocks. D) Average distal and proximal GRAB_CA_ peak heights, normalized to maximum peak height and binned into 5 blocks across training. E) Average CS+ traces across all rats for the first, middle, and final third of training F) Average CS+ and CS-traces on the final session of training for all rats. Black bars indicate where CS+ DA signal is higher than CS-DA (95% confidence intervals do not overlap).

A CS- (tone or white noise) predicted no reward. During early training, photometrically recorded GRAB_DA_ signals tended to peak following the proximal cue (syringe pump onset; Figure 2C-E). As training progressed, peak GRAB_DA_ signals migrated to the distal cue (CS+ onset; Figure 2C-E). Analyzing GRAB_DA_ peak height across training blocks reveals an interaction between Training block and Cue (proximal/distal) (F(1.921, 13.45)=5.698, p=0.0169, 2-Way ANOVA, Fig. 2D). The increase in NAc GRAB_DA_ signaling to the distal cue across training was accompanied by and strongly correlated with increased nosepoking in the reward port during the period between CS+ and reward delivery (Fig S2E,F). By the end of training, CS+ trials had higher peak GRAB_DA_ signals for distal cues compared to CS-trials, and peak heights were higher for distal than proximal peaks (Figure 2F). Migration of NAc dopamine signals to the most distal predictors of reward and reduction in reward and proximal cue dopamine signals is consistent with prior observations during Pavlovian learning[10].

### DOI restores NAc DA signals to predictable, proximal water cues and rewards

After learning was established, we gave rats a low, medium, or high cumulative dose of psychedelic 5HT_2A_ agonist, DOI, or vehicle injections, prior to water reinforced sessions consisting of ‘Expected’ and ‘Unexpected’ trial types. For ‘Expected’ trials, the water pump onset was preceded by a CS+, whereas in ‘Unexpected’ trials no CS+ was present and the onset of the water pump was unpredictable. Dopamine traces and behavioral traces for each dose and trial type can be found in Figure S2, with the primary results summarized below and in Figure 3.

**Figure 3:**
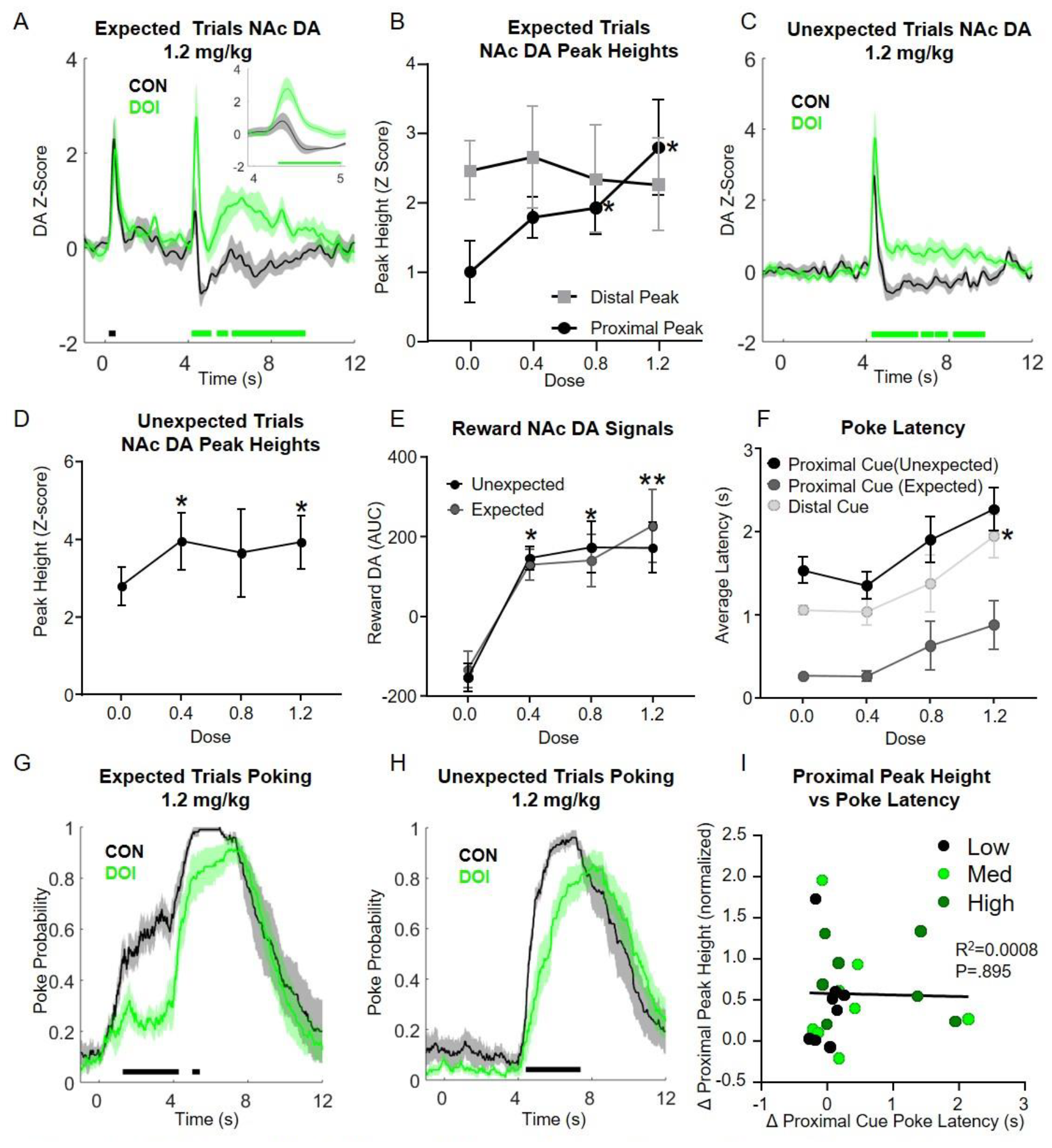
DOI Effects on NAc GRAB_DA_ and behavior during auditory-cued water delivery: A) Average traces from expected trials (trials containing a distal cue) at 1.2 mg/kg DOI. Inset: Zoomed image of proximal cue epoch B) Average GRAB_DA_ peak heights for proximal and distal cues during expected trials. C) Average GRAB_DA_ traces for unexpected trials (no distal cue) at 1.2mg/kg DOI D) Unexpected trial GRAB_DA_ peak heights. E) Reward period (3 s following pump off) DA traces across dose. F) Latency to poke in reward well by cue and dose. G) Average poke probabilities for expected trials at 1.2 mg/kg DOI. H) Average poke probabilities for unexpected trials for 1.2 mg/kg DOI. I) Linear correlation across individual sessions between change in poke latency (relative to control sessions) and change in GRABDA proximal peak heights (relative to control sessions and normalized by control distal peak heights). *All traces*: Shading indicates SEM. Green and black bars above the x-axis indicate periods of significant differences between control and DOI traces. *Line graphs*: ‘*’ and ‘**’ indicate post-hoc tests with p<0.05, and p<0.01, respectively (Dunnett‘s).

During ‘Expected’ trials, we found DOI dose-dependently increased NAc GRAB_DA_ signals to fully predictable proximal cues, without affecting GRAB_DA_ signals to distal cues (Fig 3A-B, Cue x Dose F(2.26, 15.79) = 3.997, p = 0.035; proximal post-hoc tests: 0.0 vs. 0.4 dose, p=0.0507; 0.0 vs. 0.8 dose, p=0.0428; 0.0 vs. 1.2 dose, p=0.0211, Dunnett’s multiple comparison test). We also observed small increases in GRAB_DA_ peak heights to proximal cues on ‘Unexpected’ trials, when these cues were not predictable (Figure 3C-D; 0 vs. 0.4 dose, p=0.034; 0.0 vs. 0.8 dose, p=0.54; 0.0 vs. 1.2 dose, p=0.033, Dunnett’s). For both trial types, we observed consistent, large increases in reward associated GRAB_DA_ signals that were near maximal even at the lowest dose of DOI (Fig. 3E). Concurrent with DOI-induced GRAB_DA_ signal changes, we observed dose-dependent increases in latency to enter the reward well following cues for both trial types (Fig. 3F; main effect of dose: F(1.52, 10.64)=6.669, p=0.0177). Post-hoc testing showed that at the highest DOI dose, the latency to enter the well after the distal cue was significantly increased (0.0 vs. 1.2, p=0.019, Dunnett’s). Correspondingly, for both expected and unexpected trial types we observed reduced probability of reward well occupancy for a period following cue onset (Fig. 3 G,H). Notably, there was no relationship between poke latency (relative to saline levels) and NAc GRAB_DA_ proximal peak heights across individuals (Fig. 3I), because some rats with large increases in proximal DA peaks did not exhibit increased poke latencies. This suggests that the differences in response latencies in the psychedelic state are not driving changes in NAc DA signaling.

### DOI similarly restores NAc DA signals to both food and water predictive proximal cues

Next, we sought to determine if psychedelic-induced increases in phasic DA are similar for water and food rewards that undergo opposite value shifts in the psychedelic state (see Fig. 1). We used sequential lever cues instead of auditory cues to determine the generality of DOI effects on NAc DA across cue modality and determine DOI effects on sign-tracking, a lever directed approach behavior that is associated with rigid associative learning[35,36]. We presented sequential lever cues by inserting and retracting the lever twice prior to reward delivery to examine whether cue predictability or temporal proximity to reward influenced NAc DA signaling. With this design, only the first lever cue presentation is surprising, which allows us to test the effects of DOI on multiple predictable proximal cue presentations of different identities (lever and pump/food hopper).

As in the prior experiment, we observed a dose-dependent increase in proximal cue associated GRAB_DA_ signals on DOI, and this effect was significant for both water and food reward, with no interaction between Dose and Reinforcer type (Fig. 4A-C; Main effect of Dose, F(1.586,9.518) =17.15, p=0.001, *water post-hoc tests*: 0.0 vs. 0.5 dose, p=0.003; 0.0 vs. 1.0 dose, p=0.003; *food post-hoc tests*: 0.0 vs. 0.5 dose, p=0.7227, 0.0 vs. 1.0 dose, p=0.0369, Dunnett’s multiple comparison test). We also observed dose-dependent increases in reward-associated GRAB_DA_ signals for both water and food reward that reached significance in post-hoc tests at the 1.0 mg/kg dose (Fig. 4D; main effect of Dose, F(1.226,7.353) =12.65, p=0.0069, *water post-hoc tests*: 0.0 vs. 0.5 dose, p=0.067; 0.0 vs. 1.0 dose, p=0.0039; *food post-hoc tests*: 0.0 vs. 0.5 dose, p=0.1063, 0.0 vs. 1.0 dose, p=0.0131, Dunnett’s multiple comparison test). For distal-cue associated GRAB_DA_ signals, overall there was a dose-dependent reduction in distal cue height (Fig. 4E), with a significant interaction between Dose and Reinforcer type, (Fig. 4E, main effect of Dose, F(1.141,6.844)=6.458, p=0.0367, Dose X Reinforcer type interaction, F(1.287,7.722)=12.63, p=0.006), indicating a differential dose response on NAc DA distal cue signaling for the two reinforcer types. While post-hoc tests indicated that the highest dose tested (1.0mg/kg) tended to reduce distal GRAB_DA_ peaks for food (*Food Distal*: 0.0 vs 1.0 dose, p=0.0576; *Water Distal*: 0.0 vs 1.0 dose, p=0.1795, Dunnett’s multiple testing correction), this effect did not reach significance.

**Fig 4:**
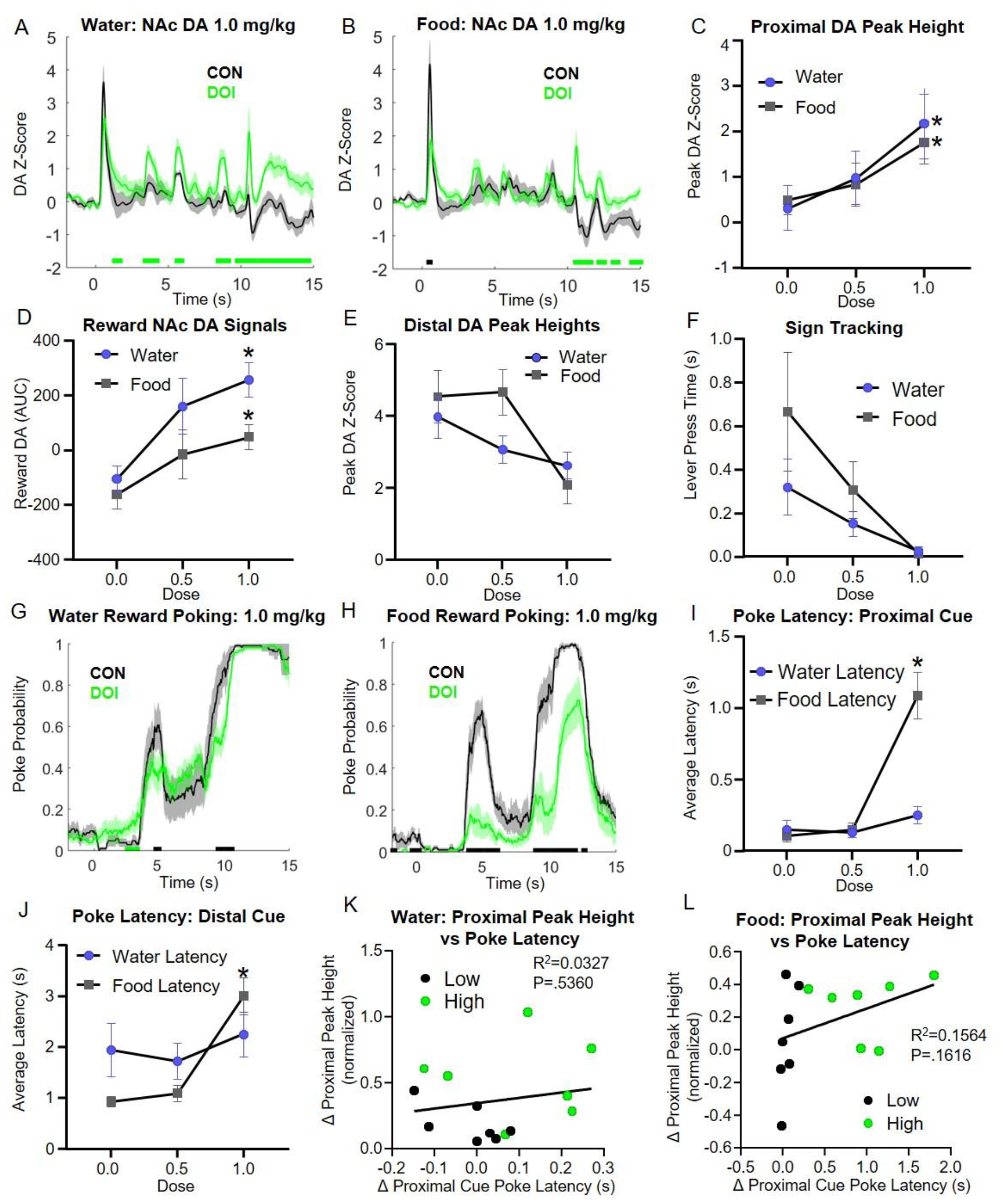
DOI Effects on NAc GRAB_DA_ and behavior during lever-cued water and food delivery: A-B) Average GRAB_DA_ traces during water **(A)** and food (B) conditions at 1.0 mg/kg. C) Proximal cue average GRAB_DA_ peak heights D) Average reward period AUC (3 s). E) Distal cue average GRABoA peak heights F) Average lever press time per trial. G-H) Average poking traces for water (G) and food (H) conditions at 1.0 mg/kg. I-J) Latency between proximal (I) and distal (J) cue and reward well poke. K-L) Linear correlation across individual DOI sessions between change in poke latency (relative to control sessions) and change in GRABoA proximal peak heights (relative to control sessions and normalized to control distal peak heights) for water (K) and food (L) conditions. *All traces:* Shading indicates SEM. Green and black bars above the x-axis indicate periods of significance between traces. *Line graphs:* ‘*’ and ‘**’ indicate post-hoc tests with p<0.05, and p<0.01, respectively (Dunnett’s).

Behaviorally, DOI dose-dependently reduced sign-tracking (Pavlovian lever pressing) for both reinforcers similarly (Fig. 4F), consistent with its effects to reduce effortful behavior across reinforcers (Fig. 1) and previous reports that DOI reduces responding for conditioned reinforcers[37]. In contrast, DOI produced markedly differential effects on poking behavior depending on the reinforcer (Fig. 4G-J). Following proximal cues, DOI increased the latency to enter the reward well for food, but not water (Fig. 4I; Dose X Reinforcer type interaction, F(1.287,7.722) = 12.63, p=0.006, *Food Proximal Latency:* 0.0 vs. 1.0 dose, p=0.003; *Water Proximal Latency:* 0.0 vs. 1.0 dose, p=0.215; Dunnett’s multiple comparisons). Similarly, for distal cues, DOI increased the latency to enter the reward well for food, but not water (Fig. 4J; Dose X Reinforcer type interaction, F(1.663,9.979) = 25.27, p=0.0002, *Food Distal Latency:* 0.0 vs. 1.0 dose, p=0.0017; *Water Distal Latency:* 0.0 vs. 1.0 dose, p=0.3959; Dunnett’s multiple comparisons).

While differences in the effects of DOI on food and water poking latencies likely reflect differences in motivation for the respective rewards, response latencies do not account for the increases observed in proximal cue associated GRAB_DA_ signals. Correlation analyses between individuals’ poking latency differences between treatments and proximal GRAB_DA_ signal differences between treatments reveal no significant relationships between these factors (Fig. 4K, L).

## Discussion

Here, we determined that the psychedelic drug, DOI, increases reward and proximal cue NAc DA signaling, despite those events being fully predictable, in Pavlovian tasks using different cue modalities and reward types. As learning progressed, DA responses were progressively inhibited to reward consumption and to fully predictable proximal reward cues in our study, replicating established results[8,10]. We show that DOI bidirectionally affects the value of food and water rewards, while DOI increases DA to proximal reward cues associated with both rewards, suggesting that changes in reward value are unlikely to explain the observed increases in DA signaling in the psychedelic state. Elevation of DA signaling to predictable proximal cues during the psychedelic state resembles prediction error signals to these stimuli observed in earlier learning stages (eg., Fig 2E) and may reflect increased error signaling even to well established associations.

Psychedelics produce a variety of behavioral disruptions that could affect NAc DA release. DOI produces hypolocomotion in rats[38], DOM and LSD increase pausing in operant responding for food[39], and DOI reduces motivation to work for rewards like food and opioids in behavioral economics tasks in rats[30], as does DOM in monkeys[40]. In the present study, we compare motivation for food and water in a behavioral economics task in the psychedelic state, finding that while food is devalued, water increases in value. We also find that as price increases, work output decreases more quickly in the psychedelic state, irrespective of the value of the reward at low prices. These data suggest that as work demands increase, motor output may become more laborious in the psychedelic state. Consistently, in Pavlovian experiments, rats tend to be slower to approach the reward well in the psychedelic state at higher doses. However, analysis of individual subjects demonstrated that many rats exhibited little or no changes in latency to approach the reward well with DOI treatment, yet exhibited large increases in DA associated with the predictable, proximal reward cue for food and water. Across individuals, there were no significant relationships between psychedelic-induced changes in approach latencies and NAc DA responses. However, we cannot rule out the possibility that behaviors we did not measure might correlate with the DOI-induced increases in NAc DA signaling observed here.

NAc DA is canonically associated with RPE[41,42], and its release decreases with the predictability of reward associated stimuli. One interpretation of the data is that the prediction (i.e., anticipation) of reward is disrupted by psychedelics. This result could be related to deficits in working memory in the psychedelic state[43,44] or difficulty in estimating temporal intervals[44,45], and this interpretation is also consistent with the view that psychedelics relax the strength of priors[23]. Another possibility is that reward prediction error signaling itself is enhanced - despite retained anticipation of the reward, *per se*. This interpretation is consistent with the observation that psychedelics can imbue ordinary stimuli with the sensation of novelty[46] (which DA is known to encode), as well as theories that posit enhancement of prediction error signaling as a core attribute of the psychedelic state[24].

NAc DA is associated with other functions besides RPE, such as the encoding of incentive salience[22], perceived salience[21], motivation[47], and costs[47–50]. An interpretation of the increased DA to proximal cues within these frameworks suggests that psychedelics may increase the salience and/or motivating aspects of proximal cues and rewards, while potentially decreasing the salience of the distal CS, evidenced by the tendency of DOI to reduce sign-tracking (associated with incentive salience) and distal cue DA - though we note DA was not consistently decreased to the distal CS for water (see Fig 3&4) across experiments. A reduction in distal cue salience is consistent with pervious work showing that DOI decreased conditioned responding to water paired stimuli, without reducing responding for water[37]. Because stimuli salience is comprised of multiple factors, including predictability, novelty, intensity, and temporally discounted value - future experiments will be required to disambiguate between these non-mutually exclusive possibilities as factors influenced by psychedelics.

As mentioned, some theoretical accounts of psychedelic action posit disruptions in predictive coding to be fundamental mechanisms by which psychedelics produce many of their subjective effects[23,24]. Two human studies show that surprising sensory stimuli produce altered EEG/MEG responses after psychedelics[51,52], though two others showed null results[53,54]. With respect to RPE specifically, one EEG study using low doses of LSD[25] and another human behavioral study of reinforcement learning[26] support the notion that psychedelics may amplify RPE processing. The results reported here further suggest that reward prediction error signaling may be enhanced by psychedelics, though more work is necessary to extend this work to sensory modalities of predictive processing.

Currently, psychedelics are under intense clinical study for varied mental health conditions including depression and drug addiction, however, a lack of mechanistic clarity on how psychedelics work is a hindrance for maximizing benefits[55]. For instance, many authors have emphasized the importance of preparation and context (‘set and setting’) in the therapeutic response(see[56] for overview), and some have written that psychedelics may function as non-specific amplifiers of the placebo response or synergize with placebo or expectancy effects[57,58]. Others have emphasized neuroplastic actions of psychedelics on dendritic structure as likely therapeutic mechanisms[59], though psychedelic-induced plasticity may be studied at several levels of inquiry from synapses to circuit level plasticity mechanisms, such as those engaged by DA neurotransmission. As DA signaling is necessary for associative learning and behavioral conditioning, further work linking psychedelic effects on DA to learning may yield additional insight into psychedelic mechanisms for producing lasting behavioral changes.

