## Supplemental Figures for "The psychedelic, DOI, increases dopamine release in nucleus accumbens to predictable rewards and reward cues"

### Slide 1
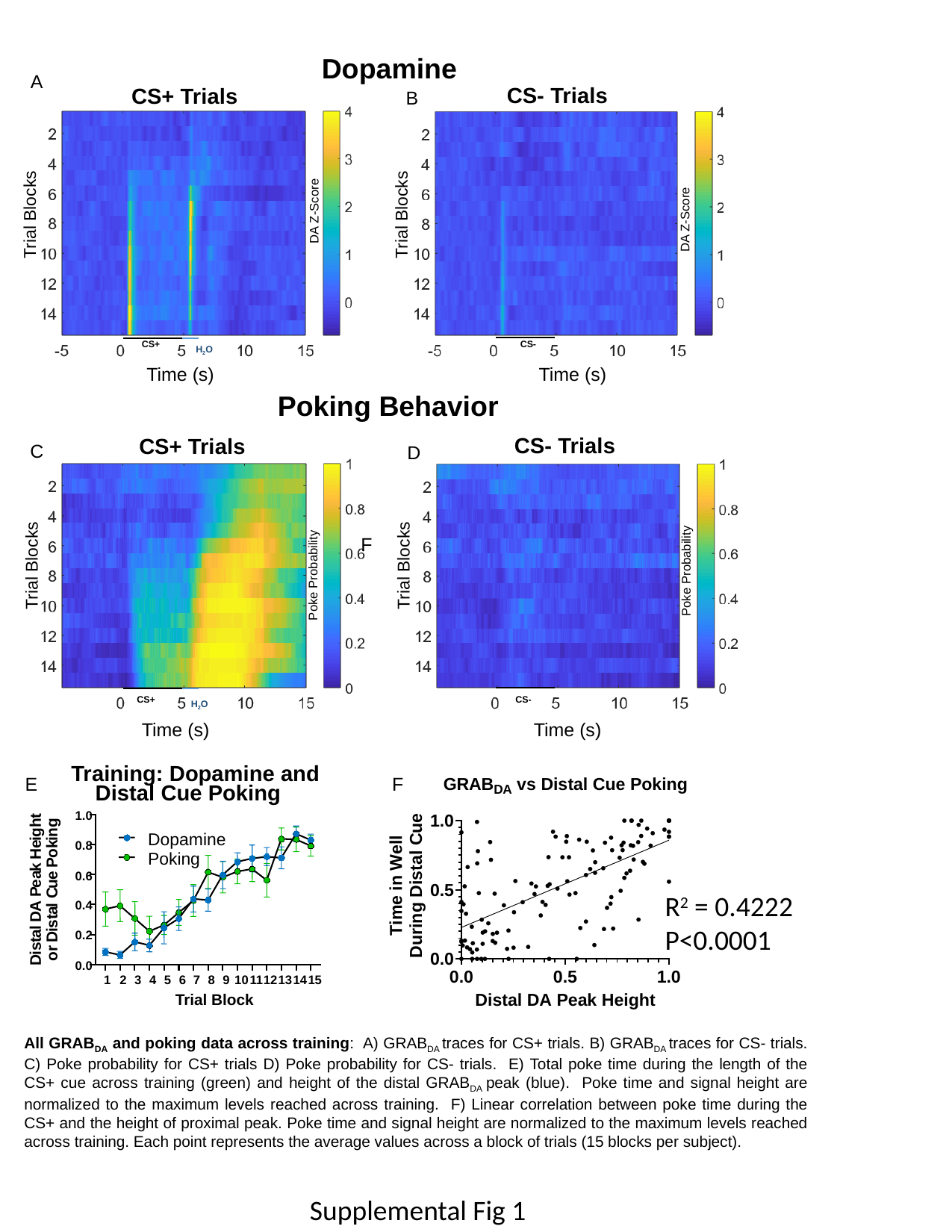

Dopamine
A
CS- Trials
CS+ Trials
B
Trial Blocks
Trial Blocks
DA Z-Score
DA Z-Score
 CS+
 CS-
 H_2_O
Time (s)
Time (s)
Poking Behavior
CS- Trials
CS+ Trials
C
D
F
Trial Blocks
Poke Probability
Poke Probability
Trial Blocks
 CS+
 CS-
 H_2_O
Time (s)
Time (s)
Training: Dopamine and
E
F
Distal Cue Poking
t
1.0
h
g
g
n
i
i
Dopamine
e
k
0.8
o
H
Poking
P
k
a
e
0.6
e
u
P
C
R^2^ = 0.4222
P<0.0001
l
A
0.4
a
D
t
s
l
i
a
D
0.2
t
s
r
i
o
D
0.0
1
2
3
4
5
6
7
8
9
10
11
12
13
14
15
Trial Block
All GRAB_DA_ and poking data across training: A) GRAB_DA_ traces for CS+ trials. B) GRAB_DA_ traces for CS- trials. C) Poke probability for CS+ trials D) Poke probability for CS- trials. E) Total poke time during the length of the CS+ cue across training (green) and height of the distal GRAB_DA_ peak (blue). Poke time and signal height are normalized to the maximum levels reached across training. F) Linear correlation between poke time during the CS+ and the height of proximal peak. Poke time and signal height are normalized to the maximum levels reached across training. Each point represents the average values across a block of trials (15 blocks per subject).
Supplemental Fig 1

### Slide 2
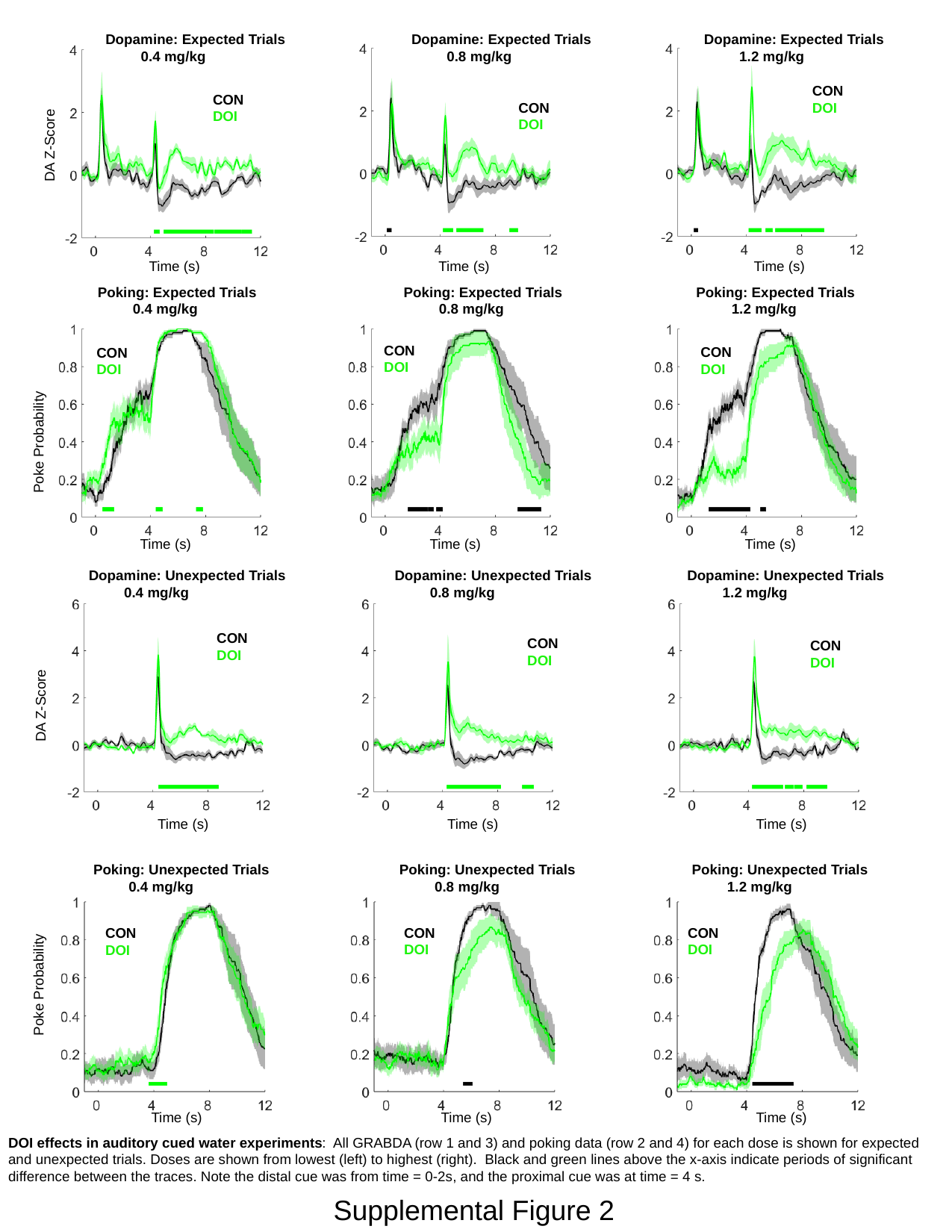

Dopamine: Expected Trials
 0.4 mg/kg
Dopamine: Expected Trials
 0.8 mg/kg
Dopamine: Expected Trials
 1.2 mg/kg
CON
DOI
CON
DOI
CON
DOI
DA Z-Score
Time (s)
Time (s)
Time (s)
Poking: Expected Trials
 0.4 mg/kg
Poking: Expected Trials
 0.8 mg/kg
Poking: Expected Trials
 1.2 mg/kg
CON
DOI
CON
DOI
CON
DOI
Poke Probability
Time (s)
Time (s)
Time (s)
Dopamine: Unexpected Trials
 0.4 mg/kg
Dopamine: Unexpected Trials
 0.8 mg/kg
Dopamine: Unexpected Trials
 1.2 mg/kg
CON
DOI
CON
DOI
CON
DOI
DA Z-Score
Time (s)
Time (s)
Time (s)
Poking: Unexpected Trials
 0.4 mg/kg
Poking: Unexpected Trials
 0.8 mg/kg
Poking: Unexpected Trials
 1.2 mg/kg
Poke Probability
CON
DOI
CON
DOI
CON
DOI
Time (s)
Time (s)
Time (s)
DOI effects in auditory cued water experiments: All GRABDA (row 1 and 3) and poking data (row 2 and 4) for each dose is shown for expected and unexpected trials. Doses are shown from lowest (left) to highest (right). Black and green lines above the x-axis indicate periods of significant difference between the traces. Note the distal cue was from time = 0-2s, and the proximal cue was at time = 4 s.
Supplemental Figure 2

### Slide 3
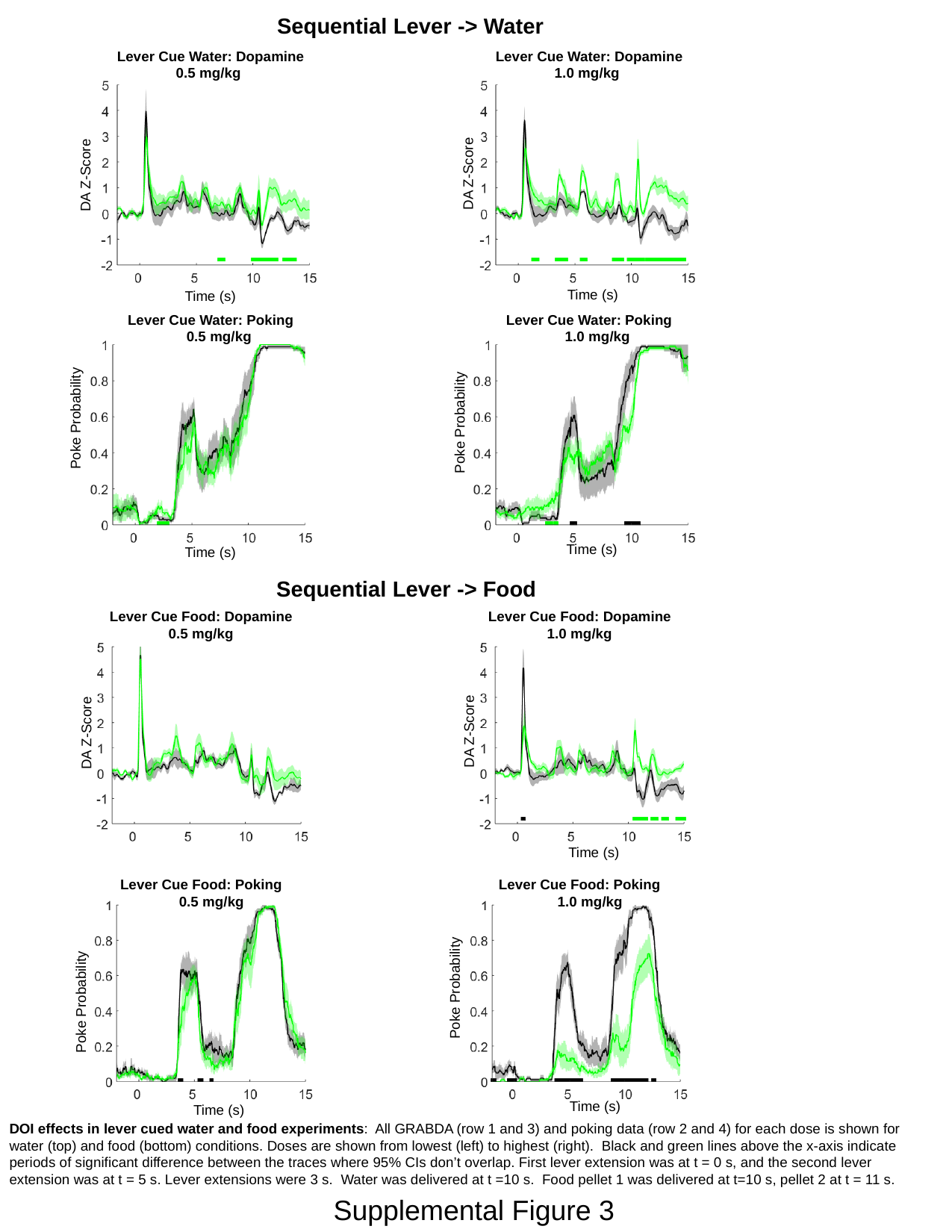

Sequential Lever -> Water
Lever Cue Water: Dopamine
 0.5 mg/kg
Lever Cue Water: Dopamine
 1.0 mg/kg
DA Z-Score
DA Z-Score
Time (s)
Time (s)
Lever Cue Water: Poking
 0.5 mg/kg
Lever Cue Water: Poking
 1.0 mg/kg
Poke Probability
Poke Probability
Time (s)
Time (s)
Sequential Lever -> Food
Lever Cue Food: Dopamine
 0.5 mg/kg
Lever Cue Food: Dopamine
 1.0 mg/kg
DA Z-Score
DA Z-Score
Time (s)
Lever Cue Food: Poking
 0.5 mg/kg
Lever Cue Food: Poking
 1.0 mg/kg
Poke Probability
Poke Probability
Time (s)
Time (s)
DOI effects in lever cued water and food experiments: All GRABDA (row 1 and 3) and poking data (row 2 and 4) for each dose is shown for water (top) and food (bottom) conditions. Doses are shown from lowest (left) to highest (right). Black and green lines above the x-axis indicate periods of significant difference between the traces where 95% CIs don’t overlap. First lever extension was at t = 0 s, and the second lever extension was at t = 5 s. Lever extensions were 3 s. Water was delivered at t =10 s. Food pellet 1 was delivered at t=10 s, pellet 2 at t = 11 s.
Supplemental Figure 3
